# Transfection of the free-living alga *Chromera velia* enables direct comparisons with its parasitic apicomplexan relative, *Toxoplasma gondii*

**DOI:** 10.1101/2025.08.26.672290

**Authors:** Isadonna F. Tengganu, Ke Hu

## Abstract

*Chromera velia* is a photosynthetic, free-living alga that is closely related to the apicomplexans, a phylum of intracellular parasites responsible for many devastating diseases, including malaria, cryptosporidiosis, and toxoplasmosis. With molecular and cellular landmarks that are clearly related to but distinguishable from those found in apicomplexan parasites, *Chromera* provides a fantastic opportunity to investigate the evolutionary origin of the structures and processes needed for intracellular parasitism. However, tools for defining localization and functions of gene products do not exist for *Chromera*, which creates a major bottleneck for exploring its biology. Here we report two major advances in exploring the cell biology of this free-living relative of a large group of intracellular parasites: 1) successful cell transformation and 2) the implementation of expansion microscopy. The initial analysis enabled by these tools generated new insights into subcellular organization in different life stages of *Chromera.* These new developments boost the potential of *Chromera* as a model system for understanding the evolution of parasitism in apicomplexans.

## Introduction

The phylum Apicomplexa has ∼ 6,000 known species of intracellular parasites [1]. This is the only large clade of eukaryotic parasites that live inside other eukaryotic cells. Many apicomplexans cause devastating infectious diseases, including malaria, cryptosporidiosis, and toxoplasmosis. Exploring the evolution of the intracellular parasitic lifestyle of these organisms will generate insights into how a cell establishes a niche inside another cell, and is also a rich source for identifying features critical for parasitism. Although we cannot travel back hundreds of millions of years to watch how free-living cells became parasites, examining and comparing the organization and functions of related structures in the parasites and their free-living relatives offers a great opportunity to determine the cellular activities that are ancestral to parasitic behaviors and how they evolved to support parasitism. *Chromera velia, a* photosynthetic, free-living relative of the apicomplexans, provides such an entry point [2]. With molecular and cellular landmarks that are clearly related to but distinguishable from those found in apicomplexan parasites, *Chromera* offers a fantastic opportunity to investigate the evolutionary origin of structures and processes needed for intracellular parasitism [2–9]. *Chromera* also bridges the apicomplexans and dinoflagellates. Those two sister clades in the superphylum Alveolata diverged more than 400 million years ago [10]. *Chromera* has readily recognizable structural and molecular features from both clades [3, 7, 11]. The phenotypic variations in these related lineages are outcomes generated from an experiment conducted in nature over several hundred million years. In the differences among these outcomes lie the answers to basic biological questions as well as the information needed to design practical solutions to some current problems in applied parasitology. Of fundamental interest are these questions: What are the requirements–molecular, cellular, and structural–for an organism to adopt and succeed in a free-living versus a parasitic lifestyle? Can we use them to define, at the molecular level, how the transition between these two life-styles occurred?

*Chromera velia* was discovered from environmental sampling of stony corals in Sydney, Australia [2]. It is straightforward to maintain and inexpensive to culture as a free-living alga. Phylogenetic and morphological analyses revealed that it is a close relative of the apicomplexans, and belongs to the chromerid lineage together with another free-living alga *Vitrella brassicaformis* [2–7]. Complete sequencing of the chromerids genomes and comparative analysis revealed major gene losses in metabolism and endomembrane trafficking pathways during the evolution from the free-living proto-apicomplexan (the common ancestor of apicomplexans and chromerids) to the immediate ancestor of the apicomplexans [7]. Intriguingly, homologs of a number of cytoskeletal genes known to be involved in parasitic behaviors in apicomplexans are found in chromerids [12–15]. Ultrastructural analysis also revealed that *Chromera* have apicomplexan-like subcellular features, including a pseudoconoid that is structurally related to but distinct from the conoid, a novel tubulin-containing structure important for *Toxoplasma* invasion into its host cell [8, 9, 14, 16–21]

To illuminate the evolutionary origin of a parasitic behavior, such as invasion, there is hardly a more powerful way than determining the localization and function of the homologs of relevant genes in the free-living relatives of the intracellular parasites. This requires the ability to specifically manipulate the sequence or the expression level of the target genes. However, while localization and molecular genetic tools are well developed for a number of apicomplexan parasites, they did not exist for *Chromera.* The field has stalled for nearly two decades at this front.

The essential requirement for genome manipulation is the transformation of the target organism, *i.e.*, the uptake of foreign DNA fragments. We have now cleared this major hurdle by developing a transfection method, and, for the first time, successfully expressed NanoLuc luciferase and fluorescent proteins in *Chromera velia*. By fluorescence-tagging, we established protein markers targeted to various subcellular structures in *Chromera*, including the nucleus, the plastid, and the cortical cytoskeleton. We discovered a *Chromera velia* promoter (pCvGAPDH) that supports cross-species expression in *Toxoplasma gondii,* which revealed an intriguing mix of divergence and conservation of protein targeting in the free-living *Chromera* and the parasitic *Toxoplasma.* To compare the cellular architectures of *Chromera* and *Toxoplasma*, we implemented expansion microscopy and labeled structures containing conserved markers for *Toxoplasma* apical and basal complexes, which produced initial insights into the evolution of cellular polarity in apicomplexans.

## Results

The life cycle of *Chromera* includes both an immotile (coccoid) stage and a motile (“zoospore”, referred to as flagellates here). The coccoid cells use the energy from photosynthesis to grow and divide in an autospore. Some of the coccoid cells can differentiate into a zoosporangium to form zoospores/flagellates in response to a light-dark cycle. In a high-light environment, the flagellates emerge from the sporangium and disseminate [11, 22]. The *Chromera* cultures used for our transfection trials typically contain mostly coccoid cells with some flagellates (**Fig 1A**).

**Fig 1.**
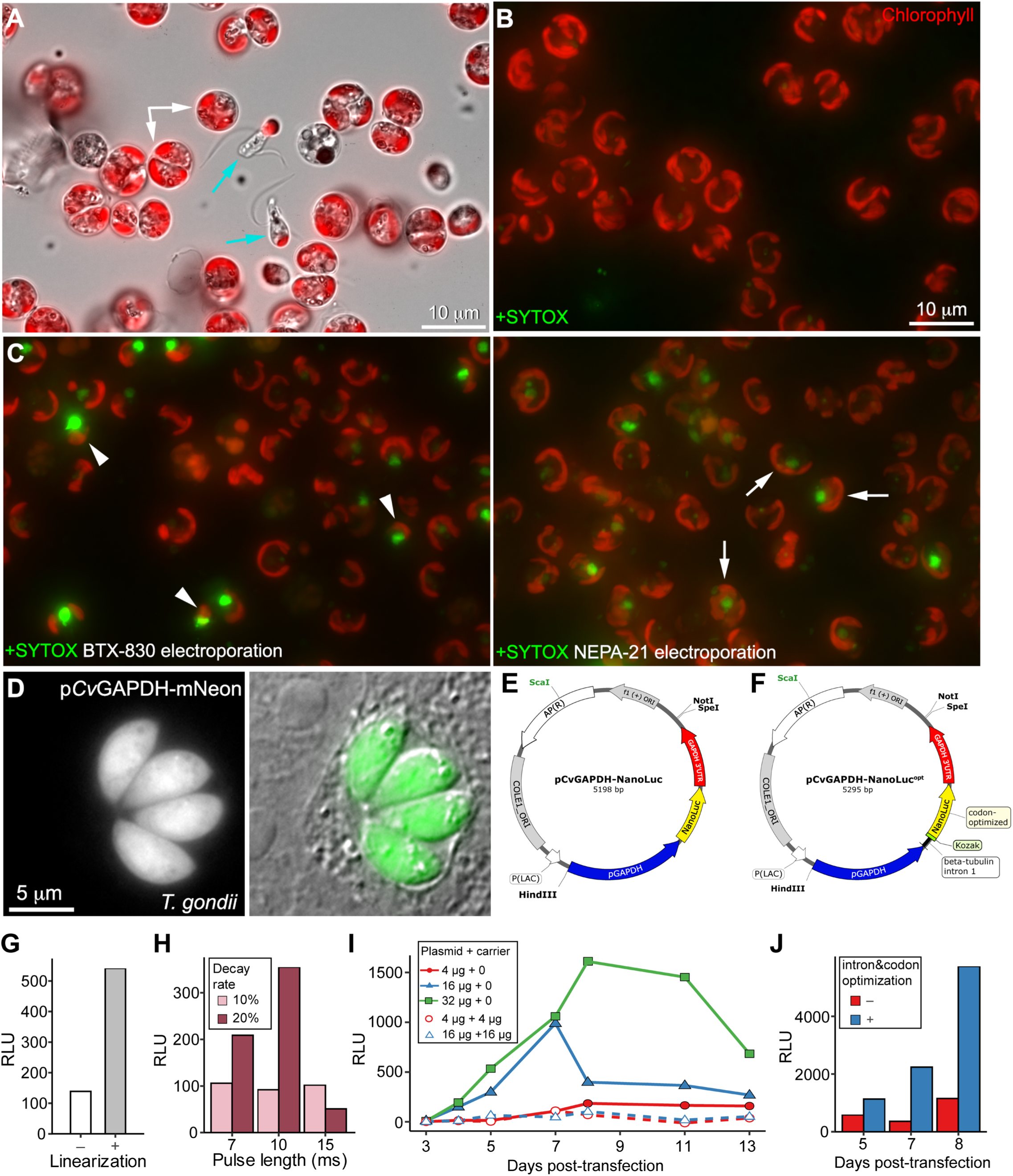
Delivery of cell-impermeant molecules into *Chromera* and expression of NanoLuc. **A.** Overlay of plastid autofluorescence (likely from chlorophyll) and transmitted light image of *Chromera velia*. White arrows: coccoid cells. Cyan arrows: zoospores (flagellates). Image contrast was adjusted to optimize display. **B.** *Chromera* coccoid cells incubated with a cell impermeant DNA dye (SYTOX^TM^Green) at 1.5 μM in sea water. Normally the plasma membrane is impermeable to SYTOX^TM^Green, as indicated by the absence of SYTOX^TM^Green fluorescence signal in the nucleus. The plastid in a coccoid typically adopts an elongated, crescent shape. **C.** Labeling of *Chromera* coccoid cells after electroporation in MEB containing 1.5 μM SYTOX^TM^Green. *Left*: BTX ECM 630 (which generates electrical pulses with exponential decay) electroporation in the presence of SYTOX^TM^Green resulted in bright labeling of the nucleus in many cells. However, the plastid often appeared to be contracted (arrowheads). *Right*: NEPA21 electroporation (which generates square-wave pulses) also generated positive SYTOX^TM^Green labeling of the nucleus but the plastid morphology is less affected (arrows). For the example in this figure, BTX ECM 630 electroporation was set at 1500 V (voltage), 50 Ω (resistance), and 25 μF (capacitance). The recorded peak voltage was 1315 V and time constant was 1.4ms. The NEPA21 electroporation was set to be two 15 ms square-wave poring pulses at 300V with 20% decay rate that delivered total poring energy of ∼ 1.65 Joule. **D.** A vacuole containing four *Toxoplasma* parasites transfected with a plasmid (pCvGAPDH-mNeonGreen) in which mNeonGreen expression is driven by the *Chromera* GAPDH promoter. **E-F**: Plasmid maps of pCvGAPDH-NanoLuc and pCvGAPDH-NanoLuc^opt^ (+intron&codon optimization) **G.** Example of comparison of effect of plasmid linearization on NanoLuc expression (7^th^ day after transfection). RLU: Relative Light Unit. All values are background-subtracted. 2.5 X 10^6^ cells were used for each electroporation. 4 μg of plasmid DNA was used with 16 µg of sheared salmon sperm DNA (SSSD) as carrier DNA. Electroporation condition: 300 V poring voltage, pulse length of 15 ms, pulse interval of 50 ms, 2 pulses, 20% decay rate with polarity switch. **H.** Example of comparison of different electroporation conditions on NanoLuc expression (7^th^ day after transfection). All experiments used 300 V poring voltage and 4 μg of *Sca*I-linearized DNA with 16 µg SSSD. **I.** Example of comparison of effect of time after transfection, amount of plasmid DNA, and SSSD carrier DNA on NanoLuc expression. For all experiments, the plasmid DNA was linearized with *Sca*I, and cells were electroporated with poring pulse at 300 V, pulse length of 10 ms, pulse interval of 50 ms, 2 pulses with 20% decay rate and polarity switch. **J.** Example of comparison of codon optimization and intron sequence on transfection using pCvGAPDH-NanoLuc (-intron&codon optimization, red) and pCvGAPDH-NanoLuc^opt^ (+intron&codon optimization, blue). For all experiments, 32 μg of plasmid DNA was linearized with NotI, and cells were electroporated with the same poring pulse parameters described in I.

Many different transformation techniques have been developed for various organisms, including electroporation, lipid-mediated gene delivery, microinjection, glass-bead-assisted gene delivery or gene-gun [23]. We focused on developing electroporation as the gene delivery method for *Chromera*, because it is scalable, amenable to optimization, and thus is the most broadly used transformation technology. Numerous cell types, including protists with different types of cell walls have been successfully transformed using this method [24–27]. There are two key steps of optimization: 1) delivery of “payload” (i.e., cell-impermeant molecules) across the plasma membrane to the intracellular environment; and 2) expression and detection of protein tracers.

### Delivery of cell-impermeant molecules

We first tested the delivery of a cell-impermeant DNA dye (SYTOX^TM^Green) by electroporation to determine the energy required to “open up” the plasma membrane. Transfection trials were carried out with user-programmable electroporators BTX ECM 630 (exponential decay, Harvard Apparatus) and NEPA21 (square wave, Bulldog Bio) apparatus. We chose these systems because the operating parameters are not hidden from the user. Both have full documentation to guide user adjustable electroporation parameters, enabling systematic tuning of energy delivered to the cells, and thus suitable for developing protocols easily adaptable by the community. BTX ECM 630 has been routinely used for *Toxoplasma* transfections [28–31]. NEPA21 has been used to successfully transfect various protists and algae [25–27]. The MAX Efficiency™ Transformation Reagent for Algae (“MEB”, A24229, Invitrogen) was used as the electroporation buffer because it was used in transformations of *Chlamydomonas*, a green alga [32]. The high impedance of the buffer (1kΩ, 50μl in 1mm gap cuvette) also prevents sparking at high voltage. Two-millimeter (mm) gap cuvettes are typically used for electroporation of eukaryotes. Because *Chromera* coccoids have a thick cell wall, we decided to use 1-mm gap cuvettes which double the strength of the electrical field at a given voltage.

Without electroporation, the plasma membrane of *Chromera* coccoid is impermeable to SYTOX^TM^Green, as indicated by the absence of SYTOX^TM^Green fluorescence signal in the nucleus (**Fig 1B**). When electroporated with BTX ECM 630 in MEB at peak voltage of ∼ 1315 V and time constant of 1.4 ms, many cells are brightly labeled in the nucleus, indicating penetration of the cell membrane (in comparison, typical peak voltage is ∼ 850 V for *Toxoplasma* transfections with BTX ECM 630 [28–31]). However, the plastid often appeared to be contracted, which is an indicator of lower viability. NEPA21 uses a different mechanism of generating electrical pulses, the poring voltage (300V) is considerably lower, but the pulse length can be up to 15 ms. Multiple pulses with a set decay rate between each pulse can be programmed. Total energy delivered can be calculated based on the voltage, impedance, pulse length, pulse numbers and decay rate. When sample impedance is at ∼ 1.37 KΩ and poring voltage at 300V, two 15 ms pulses with 20% decay rate deliver total poring energy of ∼ 1.65 Joule. At this condition, NEPA21 electroporations in MEB generated positive SYTOX^TM^Green labeling of coccoid nucleus and the plastid morphology is less affected than in BTX ECM 630 electroporations (**Fig 1C**). We therefore mainly used NEPA21 for optimizing *Chromera* transfection in the subsequent experiments.

### Selection of tracer proteins and promoters for optimization of transfections

Besides the uptake of molecules, the apparent transfection efficiency for an expression plasmid also depends upon the detection efficiency, which is affected by the level of expression. It also depends on sensitivity of detection of the protein product, which is affected by promoter strength and brightness of the tracer proteins. The combination of strong promoters and super bright tracer proteins decreases the number of DNA molecules required for yielding a detectable signal and allows the detection of protein expression even at very low transfection efficiencies, which provides the foundation for further optimization using a matrix of transfection conditions.

We reasoned that promoters for house-keeping genes known to be highly transcribed universally in eukaryotes are good candidates for the transfection trials. We started with promoters for the highly conserved GAPDH (Glyceraldehyde 3-phosphate dehydrogenase) and ribosomal protein L31. In *Toxoplasma,* the transcription levels of GAPDH and L31 are ranked as top ∼ 20% and top ∼ 1% of the entire transcriptome, respectively. ∼ 1.1 kb genomic sequence upstream of the coding sequence for *Chromera* GAPDH (Cvel_5775, VEupathDB.org) and L31 (Cvel_7598) were used as promoters to drive the expression of tracer proteins. 632 bp (GAPDH) or 499 bp (L31) downstream of the stop codon of the CDS were used as the 3’UTR. As an initial assessment, we tested the ability of these *Chromera* promoters and 3’ UTRs to support expression of the fluorescent protein mNeonGreen in *Toxoplasma.* We found that pCvGAPDH, but not pCvL31, drove robust expression of mNeonGreenFP in *Toxoplasma* (**Fig 1D**). The pCvGAPDH-driven expression in a heterologous system not only reveals the conservation of the basic transcriptional machinery but also indicates strong promoter activity. Therefore, we used pCvGAPDH in the subsequent experiments to optimize transfection in *Chromera*.

### Detection and optimization of NanoLuc expression

Even though pCvGAPDH-mNeonGreen showed robust expression in *T. gondii, Chromera* transfection trials with this plasmid yielded no positive results. When the transfection efficiency is low, scanning for cells expressing fluorescent protein is slow and tedious, which is a major barrier for testing many different transfection conditions. Ultrasensitive bulk assays that can detect transfection in a vanishingly small fraction of a large population are needed. We thus decided to use NanoLuc luciferase (NanoLuc) as the tracer molecule, because it has extremely robust enzymatic activity for generating luminescence using furimazine as the substrate [33]. It has been used for several new model systems in recent years, such as Choanoflagellate and *Crytosporidium* [34, 35]. pCvGAPDH drives robust NanoLuc expression in *Toxoplasma ∼* 24 hr after transfection (>100,000 ∼ fold above background). However, no expression could be detected in the initial many trials of transfections in *Chromera* in which we mainly focused on varying electroporation parameters. We therefore embarked on a large number of trials in which we iteratively tuned multiple parameters, including plasmid reengineering, expression time, amount of plasmid DNA, as well as the inclusion of carrier DNA. We also developed an assay buffer and sample preparation method to measure the luciferase activity of NanoLuc expressed in *Chromera* cells, which gave us a higher reading than using the standard commercial kit and protocol. The progression of optimization is far from linear, and includes iterative testing of multiple factors. Here we will just give a few examples of the conditions that we screened, which includes the factors that appear to have consistent impact on NanoLuc expression as assessed by luminescence. 2.5 X10^6^ to 5 X10^6^ cells were used in each transfection.

We first codon-optimized the NanoLuc CDS for expression in *Chromera*. We also inserted an intron from a *Chromera* β-tubulin (Cvel_33153) between the promoter and the kozak sequence, because the inclusion of introns has been shown to significantly increase gene expression in metazoa, fungi, plants, and protists, through, for example, boosting the stability of a transcript [36]. Inserting the intron before the coding sequence also helps to simplify subsequent modification of this plasmid to express different transgenes. This new NanoLuc plasmid was named pCvGAPDH-NanoLuc^opt^ (see **Fig 1E-F** for comparison between pCvGAPDH-NanoLuc and pCvGAPDH-NanoLuc^opt^).

Our initial transfections with circular pCvGAPDH-NanoLuc^opt^ did not yield significant signal after 48 hr, but very weak expression could be detected on day 5. As linearization of the plasmid has been shown to affect expression levels in other systems, we linearized the pCvGAPDH-NanoLuc^opt^ plasmid (**Fig 1G-H**) and further extended the timeline for measuring the NanoLuc activity, which showed a consistent trend of increasing total NanoLuc activity up to 7-8 days (**Fig 1I-J**). The amount of plasmid DNA and the presence or absence of carrier DNA also had an impact on NanoLuc expression (**Fig 1I**). In parallel to testing the impact of DNA preparation on transfection efficiency, we also examined the impact of electroporation conditions, with varying voltage, pulse length, and decay rate (**Fig 1H**). After the pCvGAPDH-NanoLuc^opt^ transfections yielded positive results consistently, we reengineered the pCvL31 plasmid to include the Cv-codon optimized NanoLuc gene and the pre-kozak sequence intron (pCvL31-NanoLuc^opt^). This modified pL31 plasmid then also yielded positive transfection results in *Chromera* (data not shown), thus identifying another promoter that can be used for expressing transgenes in *Chromera*.

After we pinned down the broad conditions for transfections (expression time, DNA preparation, and electroporation conditions), we were able to obtain positive NanoLuc expression (estimated by luminescence reading) without fail. However, the absolute luminescence readout varied considerably between experiments and sometimes even between replicates. The variations were likely caused by multiple factors, such as growth conditions (e.g. how long the cells have been in culture), or sample preparation for measuring luminescence (e.g. sonication in water bath to break up the cells for better NanoLuc/furimazine interaction). Because of these variations and also because our primary goal is to localize proteins by tagging with fluorescent proteins, we decided to not further fine-tune the transfection using the NanoLuc assay, but instead start testing transfections of fusion genes encoding *Chromera* proteins tagged with fluorescent proteins.

### Comparison of subcellular targeting in live Chromera and Toxoplasma by fluorescent-protein tagging and cross-species expression

We selected several target proteins that are expected to localize to different subcellular structures in *Chromera* based on the known localization of their apicomplexan homologs. For each transfection, 2.5 × 10^6^ cells were electroporated with 16 µg of linearized plasmid (double-cut with *Hind*III and *Spe*I, see Methods) without carrier DNA using the following poring pulse parameter: poring voltage 300 V, pulse length of 10 ms, pulse interval of 50 ms, 2 pulses with 20% decay rate and polarity switch. Fluorescence was typically observed 4 days after transfection and imaged between day 4 and day 6. The apparent transfection efficiency varies among constructs. For transfections that express histone H2B-mStayGold, the highest apparent transfection efficiency is ∼ 0.2% (∼20,000 total cells counted).

The *Chromera* genome contains a full set of canonical core nucleosome histones (H2A, H2B, H3, and H4) that are highly conserved with *Toxoplasma* histones. We found that a *Chromera* histone H2B (CvH2B, Cvel_19200) tagged with mStayGold [37] is specifically targeted to the nucleus of *Chromera*. The labeling of CvH2B-mStayGold is not uniform across the nucleus, which is likely due to variations in local chromatin condensation (**Fig 2A**). In addition to single cells, we also found CvH2B-mStayGold-expressing cells that recently completed nuclear division or cell division (**Fig 2B**-**C**). In one pre-cytokinesis cell, the two divided nuclei are positioned nearly symmetrically at the ends of one cell axis, while the plastid forms a complete ring (**Fig 2B**, top). In another example, cytokinesis has completed, but the new cells remain tightly bound. This cluster of cells contain four nuclei labeled by CvH2B-mStayGold, indicating the expression of the fusion protein can persist over at least two generations (**Fig 2B**). When expressed in *Toxoplasma,* CvH2B-mStayGold also highlights the nuclear chromatins, suggesting conserved nucleosome packing (**Fig 2D**).

**Fig 2.**
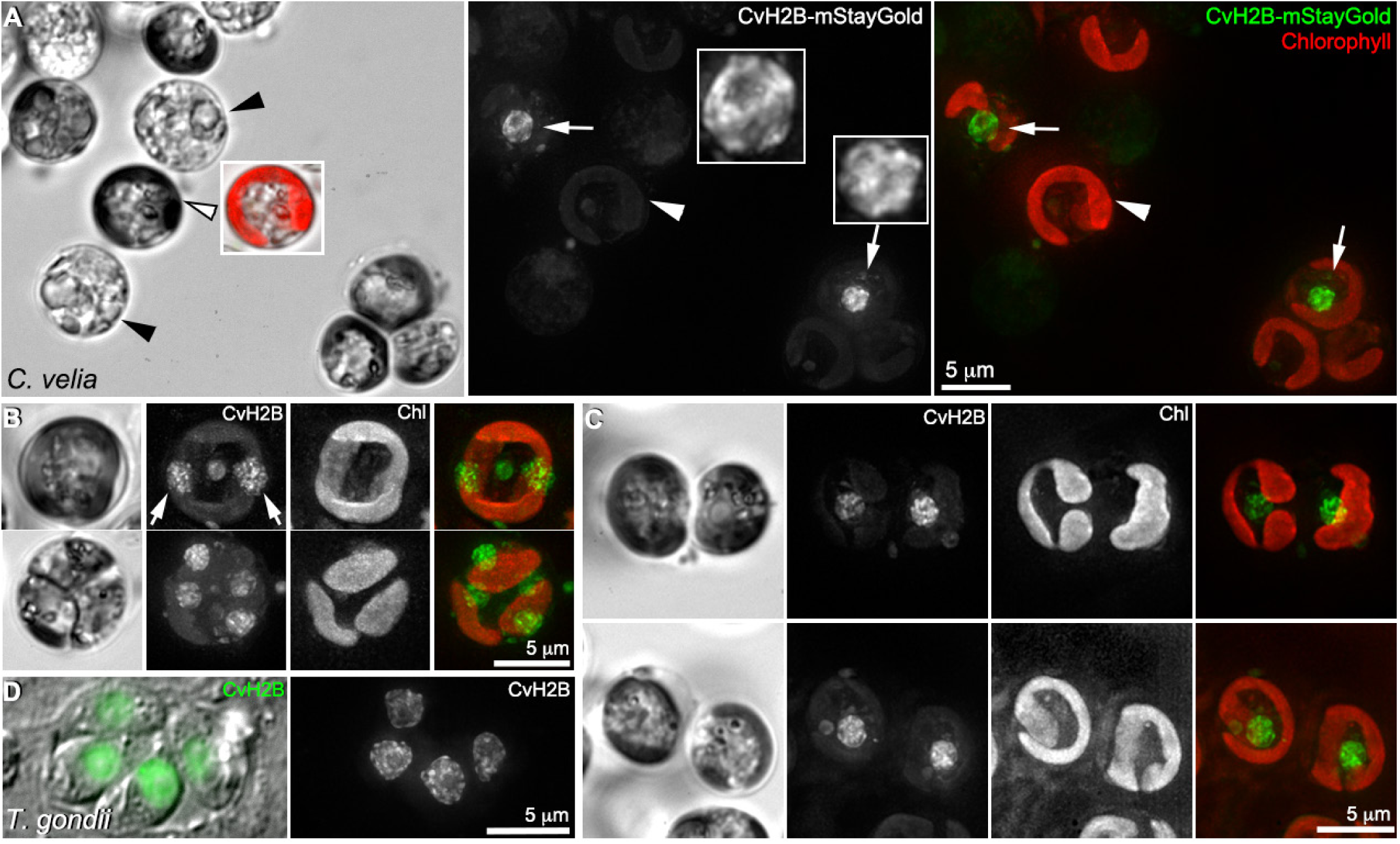
CvH2B-mStayGold highlights the nuclear chromatin in *Chromera* coccoids. **A.** A field containing multiple *Chromera* coccoids, including two that express CvH2B-mStayGold (arrows, 3X inset). White arrowheads: the pigments in the plastid mainly fluoresce in the red to far-red channel, but weak bleed-through in the green channel can also be observed. The plastid appears to be darker in the bright field image (left) due to light absorption (inset: overlay of chlorophyll fluorescence and transmitted light image). Black arrowheads: Some cells in the population appear to have lost the chlorophyll and are presumably dead. Fluorescence images are projections of stacks of deconvolved wide-field images. Image contrast was adjusted to optimize display. **B.** CvH2B-mStayGold-expressing cells that recently completed nuclear division (top) or cell division (bottom). Top panels show a pre-cytokinesis cell, in which the two divided nuclei are positioned nearly symmetrically at the ends of one cell axis (arrows). The spherical object at the center of the cell is from autofluorescence. Bottom panels show a cluster of cells that divided recently but remain tightly bound, and the cells have not yet adopted the spherical shape of a typical coccoid cell. This cluster of cells contains four nuclei labeled by CvH2B-mStayGold, indicating that the expression of the fusion protein can persist over at least two generations. Chl: chlorophyll. **C.** Pairs of *Chromera* coccoids expressing CvH2B-mStayGold. These cells, likely having divided recently, have developed a spherical shape, but remain close. **D.** A vacuole containing four *Toxoplasma* parasites transfected with pCvGAPDH-CvH2B-mStayGold. CvH2B-mStayGold is specifically targeted to the nuclear chromatin.

The discovery of a degenerated plastid in apicomplexans, including *Toxoplasma* and *Plasmodium*, was one of the earliest observations supporting the descent of these parasites from free-living organisms [38–40]. The non-photosynthetic apicomplexan plastid is critical for fatty acid synthesis, and contains many components for acyl chain synthesis and transfer, including the nuclear encoded Acyl Carrier Protein (ACP) [41–43]. In both *Toxoplasma* and *Plasmodium,* ACP contains a N-terminal bipartite targeting sequence, a signal sequence that allows the protein to enter the endomembrane system, and a transit peptide recognized by the plastid importers [41, 42]. Without the transit peptide, the protein is secreted to the parasitophorous vacuole. Without the signal peptide, the protein remains in the cytoplasm [41, 42]. *Chromera* has a strong homolog of TgACP. When tagged with mStayGold at the C-terminus, CvACP (Cvel_32060) is targeted to the plastid in *Chromera* coccoids (**Fig 3A-B**). In most cells, the fluorescence of CvACP-mStayGold completely overlaps with the signal from chlorophyll. However, we note that in some *Chromera* cells CvACP-mStayGold is targeted to the rim of the cell in addition to the plastid (**Fig 3C**). It is possible that high level of over-expression of CvACP-mStayGold overwhelms the import machinery to the plastid, which leads some fusion proteins to be secreted and trapped between the plasma membrane and coccoid cell wall. Interestingly, while CvACP contains the full targeting signal to enter the *Chromera* plastid, when expressed in *Toxoplasma,* CvACP is secreted to the parasitophorous vacuole (**Fig 3D**), indicating that the signal sequence of CvACP is recognized by the ER for the protein to enter the endomembrane pathway, but not by the plastid import machinery in the parasite, suggesting a divergence of the membrane trafficking system.

**Fig 3.**
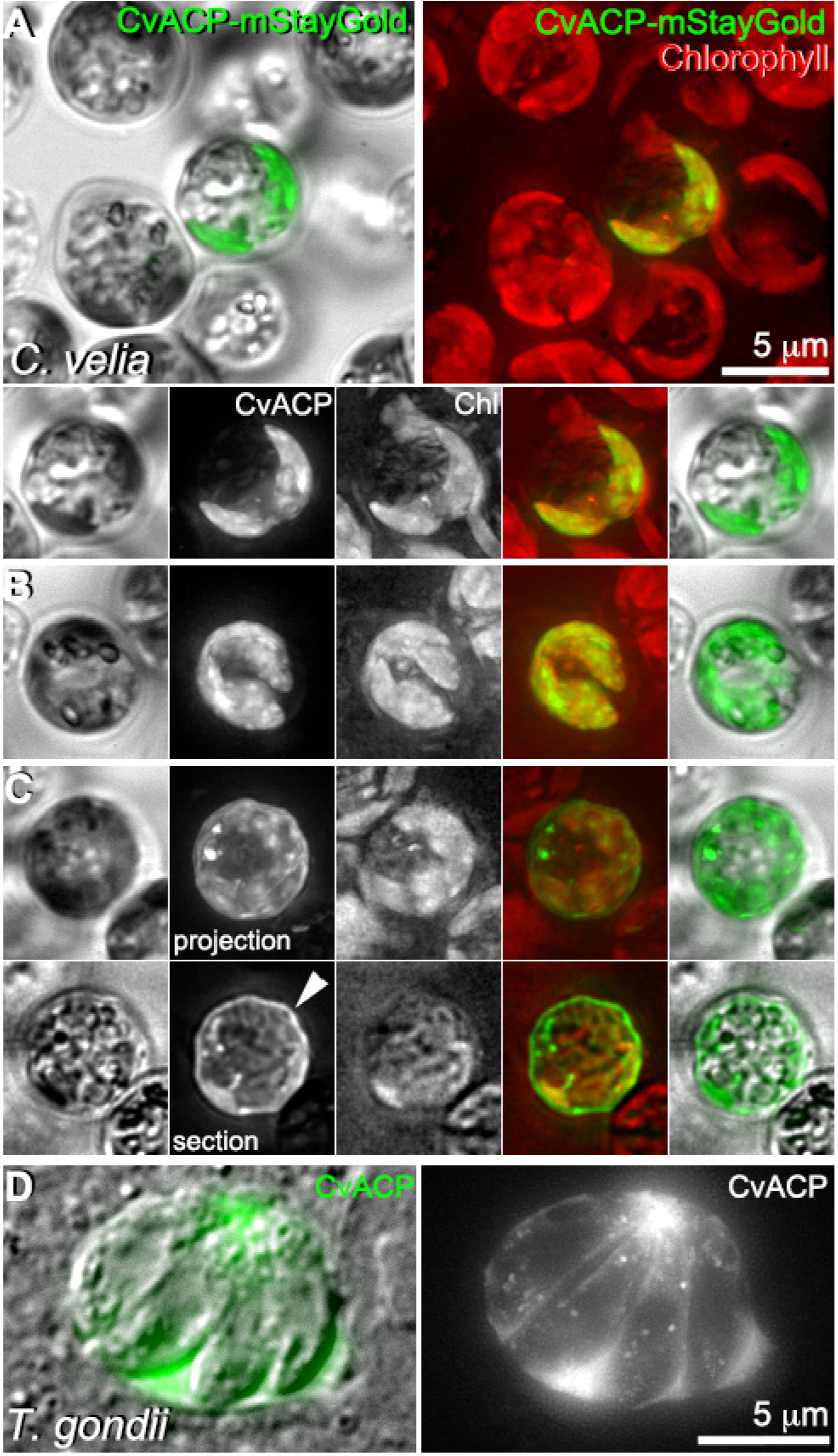
CvACP-mStayGold is targeted to the plastid in *Chromera* coccoids, but is secreted to the parasitophorous vacuole when expressed in *Toxoplasma*. **A-B.** CvACP-mStayGold (green) is targeted to the plastid in *Chromera,* as indicated by the signal overlapping with chlorophyll fluorescence (red). Chl: chlorophyll. Image contrast was adjusted to optimize display. **C.** In addition to the plastid, CvACP-mStayGold is targeted to the cell rim of some *Chromera* coccoids (arrowhead). A projection (top) and a single plane (bottom) of the fluorescence images are shown. **D.** A vacuole containing four *Toxoplasma* parasites transfected with pCvGAPDH-CvACP-mStayGold. CvACP-mStayGold (green) is secreted out of the parasites into the parasitophorous vacuole.

All apicomplexans have an intermediate-filament-like network associated with the cortex. The components of this network belong to the alveolin-family proteins that unite the three major clades of Aveolata, apicomplexans, dinoflagellates, and ciliates [44]. The alveolins associate with the alveoli, which are membrane-enclosed sacs underlying the plasma membrane. In apicomplexans, these sacs are flattened and sutured together to form the inner membrane complex, and the apicomplexan alveolins are named as IMC proteins [45–49]. We examined the localization of a *Chromera* IMC protein, Cvel_9514. Because it shows closest homology with TgIMC13 [48], we tentatively name it CvIMC13. In *Chromera,* CvIMC13 is targeted to series of cortical ring (**Fig 4A**) or arc-shaped (**Fig 4B**) structures, a pattern unknown before. When expressed in *Toxoplasma,* CvIMC13 is largely cytoplasmic, with, however, enrichment in the cortex of developing daughters (**Fig 4C**). This indicates that while CvIMC13 can be tolerated by the daughter IMC network, it cannot be efficiently retained in the IMC network in adult parasites.

**Fig 4.**
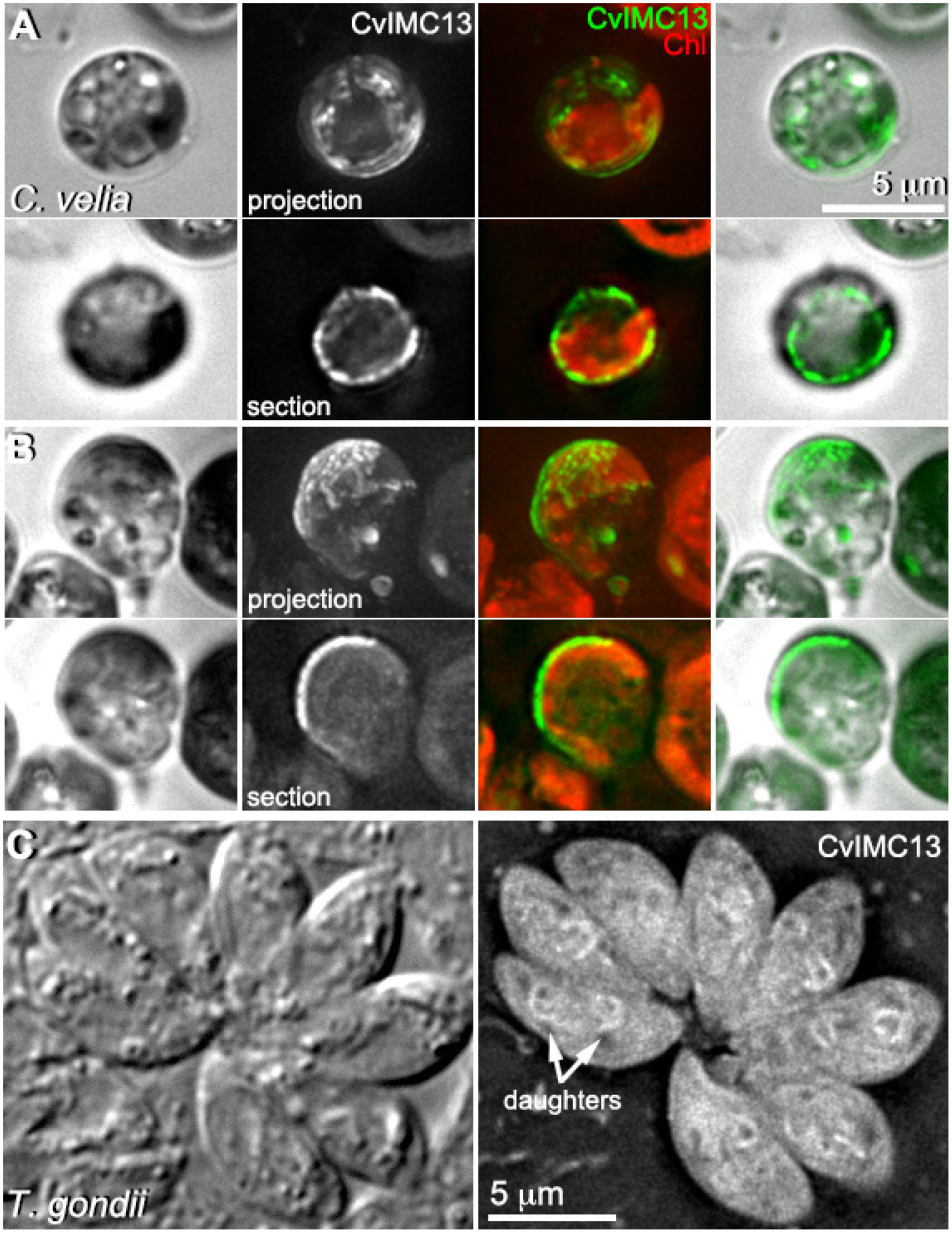
CvIMC13-mStayGold highlights cortical structures in *Chromera* coccoids. **A-B.** CvIMC13-mStayGold (green) is targeted to series of ring (A) or arc-shaped (B) structures in the cortex of *Chromera*. Chl: chlorophyll. A projection (top) and a single plane (bottom) of the fluorescence images are shown. Image contrast was adjusted to optimize display. **C.** A vacuole containing eight dividing *Toxoplasma* parasites transfected with pCvGAPDH-CvIMC13-mStayGold. CvIMC13A-mStayGold is mostly cytoplasmic, with some enrichment in the daughter cortex.

One of the most fascinating cellular architectures in apicomplexans is the cortical microtubule (MT) array. In *Toxoplasma,* the 22 cortical MTs originate from the apical polar ring and form an ultrastable rib cage [29, 50–52]. The *Chromera* genome encodes two canonical α-tubulin genes coding for identical amino acid sequences and three copies of canonical β-tubulins. Expressing the Cv-α-tubulin (Cvel_22324) with mStayGold at its N-terminus highlights fibers that cover a portion of the *Chromera* coccoid (**Fig 5A**). When expressed in *Toxoplasma,* Cv-α-tubulin-mStayGold is incorporated into multiple tubulin-containing structures (**Fig 5B**). This includes structures formed of canonical MTs, such as the cortical MT array and the spindle poles, as well as the conoid, which is formed of highly curved, non-tubular, tubulin-containing fibers [16]. The conoid is the core structure of *Toxoplasma* apical complex (**Fig 6A**). It is structurally related to the pseudoconoid in the *Chromera* flagellates, which is formed of curved-MTs [8, 9]. However, as a pseudoconoid or its precursor has not been found in the coccoid cells, it is difficult to compare the cellular architecture between the *Chromera* and the apicomplexans at this cell stage. The relatively low transfection efficiency so far has prevented us from identifying any flagellates expressing fluorescently tagged proteins.

**Fig 5.**
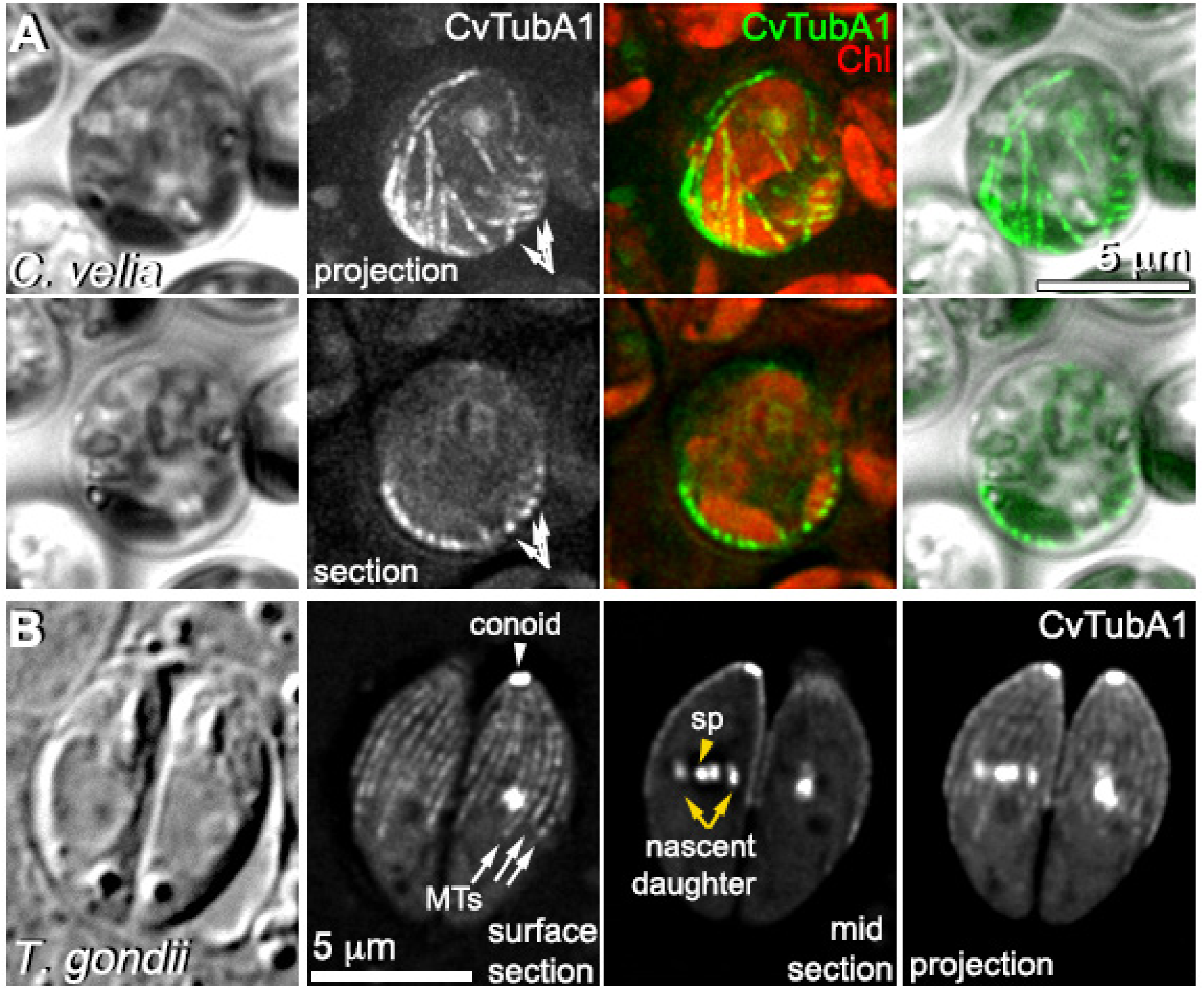
mStayGold-CvTubA1 highlights cortical fibers in *Chromera* coccoids. **A.** mStayGold-CvTubA1 highlights cortical fibers (arrows) of the *Chromera* coccoid cells. A projection (top) and a single plane (bottom) of the fluorescence images are shown. Image contrast was adjusted to optimize display. **B.** A vacuole containing two *Toxoplasma* parasites transfected with pCvGAPDH-mStayGold-CvTubA1. mStayGold-CvTubA1 is targeted to multiple tubulin containing structures, including the conoid, cortical MTs, and spindle poles (sp). The tubulin-containing cytoskeleton of nascent daughters (yellow arrows) are forming in these parasites. A surface section, a middle section, and a projection of the fluorescence images are shown.

**Fig 6.**
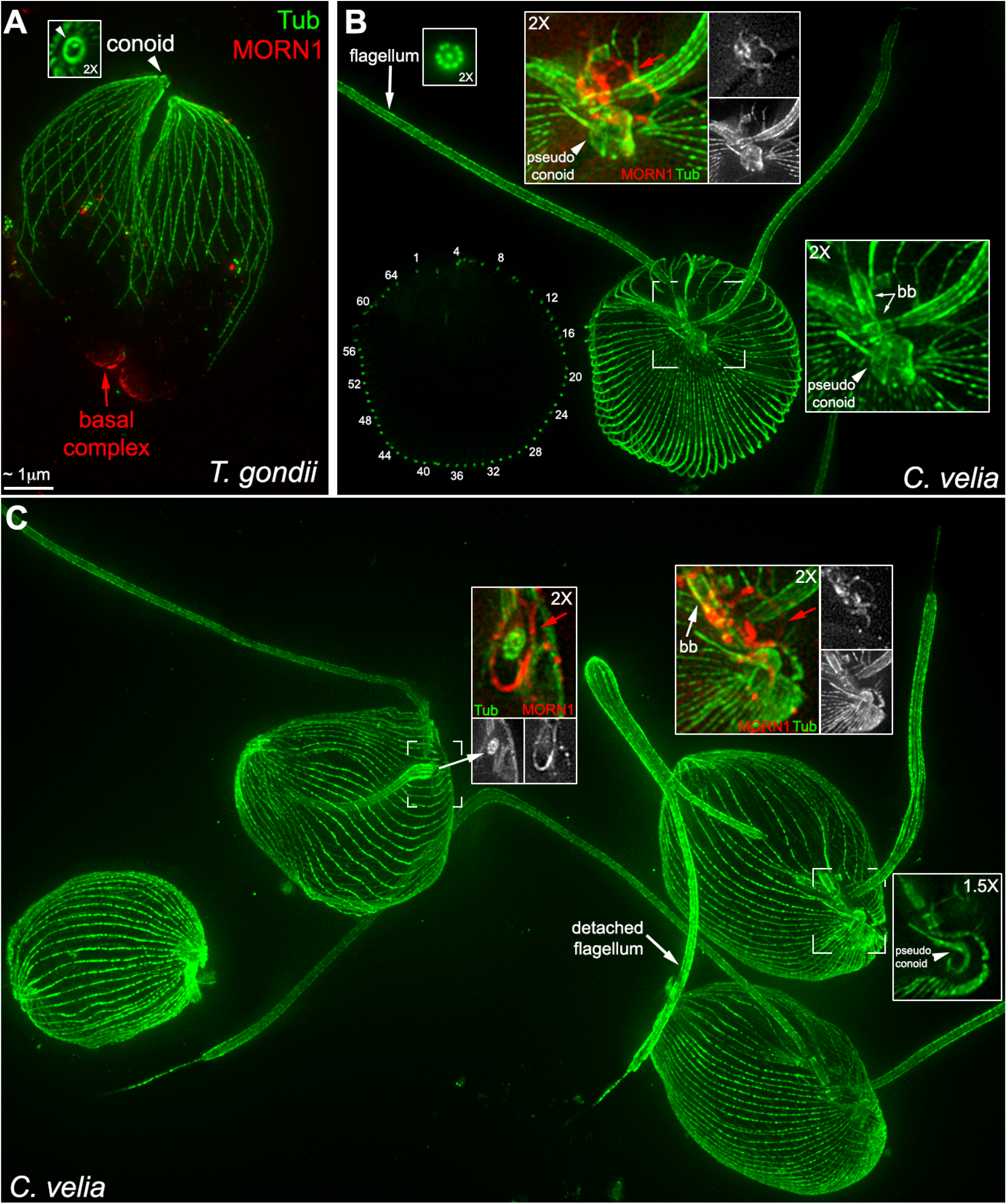
Comparison of the cytoskeletal architectures of *Toxoplasma* and *Chromera* flagellates using expansion microscopy (ExM). **A.** Projections of ExM images of two*Toxoplasma* parasites labeled with anti-tubulin (green) and anti-MORN1 (red) antibodies. The conoid (white arrowhead) and the basal complex (red arrow) mark the apical and basal poles of the parasite. The inset (2X, image from a different parasite) shows that the conoid wall is closed into a circle in the end-on view. The spot in the middle of the conoid is the intraconoid MTs. Image contrast was adjusted to optimize display. Scale bar is based on an estimated expansion ratio of ∼5.4 [52, 62]. **B.** ExM images of a *Chromera* flagellate. *Main image*: a full projection through a flagellate, in which the cortical MTs, pseudoconoid, basal bodies, and flagella are labeled with anti-tubulin antibodies. *Left*: A single section through the flagellate that shows 64 MT cross-sections around the cortex. Left inset (2X, image from a different cell): cross-section view of a flagellum, in which the 9 doublet and central pair MTs can be resolved. The middle and right insets include the region indicated by the bracket. Right inset (2X): Pseudoconoid, which is built from curved MTs, is positioned close to the two perpendicularly positioned basal bodies (bb). Middle insets include a projection of a substack of the region around the base of one of the flagella. The overlay of anti-MORN1 (red) and anti-tubulin (green) is displayed at 2X. Anti-MORN1 highlights structures that associate with the basal body and encircle the base of a flagellum (red arrow). Anti-MORN1 labeling is only shown in the projection of a substack because the signal is weak, and a full projection interferes with viewing the structures of interest. Image contrast was adjusted to optimize display. **C.** ExM images of several *Chromera* flagellates. *Main image*: a full projection of multiple flagellates labeled with anti-tubulin antibodies. Insets include the region indicated by the bracket. Left and middle insets include anti-MORN1 labeling, which consistently highlights structures that associate with the basal body (bb) and encircle the base of a flagellum (red arrows). The insets on the left include a near cross-section view of a flagellum (white arrow) positioned in the middle of the MORN1 ring (red arrow). The overlays of anti-MORN1 (red) and anti-tubulin (green) are displayed at 2X. Right inset (1.5X): The projection of a substack that shows a cross-section of the pseudoconoid wall, which is highly curved and open on one side. Contrast this with the end-on view of *Toxoplasma* conoid in the inset in panel A.

### Comparison of the cytoskeletal architectures of Chromera flagellates and Toxoplasma using expansion microscopy (ExM) suggests that the basal complex in apicomplexans might have originated from a flagellum-associated structure

To examine the cellular architecture of the *Chromera* flagellate, we used expansion microscopy to generate, for the first time, a 3-D view of its tubulin-based cytoskeleton (**Fig 6B**). The MT array of flagellated *Chromera* form a striking corset that covers the entire cell, which is distinct from the MT array in *Toxoplasma*, where there is a clear separation between the 22 cortical MTs and the basal pole, which is capped by the basal complex (**Fig 6A**). Two flagella originate from two basal bodies that juxtapose perpendicularly. The pseudoconoid is located close to the basal bodies, consistent with previous electron microscopy observations [9] (**Fig 6B**, right inset).

The cellular polarity of the apicomplexans is defined by the apical and basal complexes. The loss of a major component of the basal complex, TgMORN1, results in a significant defect in cytokinesis in *Toxoplasma* [53–56]. *Chromera* has a super-conserved homolog of MORN1 (Cvel_32330). In *Chromera* flagellates, an anti-TgMORN1 antibody [55] highlighted ring or arc shape structures encircling the base of the flagella. Some anti-MORN1 signal also associated with the basal bodies (**Fig 6B**-**C**, insets). It was proposed that the apical complex in *Toxoplasma* is evolutionarily connected with the flagellum, due to several shared components and the consistent association between the pseudoconoid and the flagellum in *Chromera* [8, 9, 52]. Our results suggests that the basal complex in apicomplexans might also have originated from a flagellum-associated structure.

## Discussion

The apicomplexans form a large phylum of intracellular pathogens. Over hundreds of millions of years, they have successfully parasitized metazoa, and cause devastating diseases in a broad range of hosts, from butterflies, sea otters, to humans [1, 57, 58]. The discovery of *Chromera* was a major milestone in apicomplexan biology, because it strongly supports the long-standing hypothesis that the apicomplexans descended from a free-living ancestor [2]. *Chromera* also provides an opportunity to directly determine how differences in molecular functions and organizations impact a free-living or parasitic lifestyle. In order to answer these questions, developing localization and molecular genetic tools is essential.

Combining optimization of multiple parameters, such as plasmid engineering, preparation, expression time, and electroporation conditions, we have now successfully expressed not only an ultrasensitive tracer protein, but also a series of *Chromera* proteins tagged with mStayGoldFP that are targeted to different subcellular locations in *Chromera.* The pCvGAPDH promoter we identified also drives robust gene expression in *Toxoplasma*, which is a convenient tool for comparing protein targeting in *Chromera* and *Toxoplasma* by cross-species expression. This comparison revealed an intriguing mix of conservation and divergence. For example, both CvH2B and TubA1 are targeted to the same subcellular locations as their *Toxoplasma* homologs. On the other hand, the *Chromera* plastid protein ACP is secreted out of the parasite into the parasitophorous vacuole, indicating the divergence of the import systems in the fully functional *Chromera* plastid and degenerated parasite plastid.

To complement live-cell imaging with fluorescent protein tagging, we have now also implemented expansion microscopy in *Chromera*. This enables the examination of the 3-D organization of the sophisticated MT cytoskeleton in *Chromera* flagellates. Comparing the relative localization of homologous components of apical and basal complexes revealed that the apical and basal poles of the apicomplexans might have arisen from structures associated with the base of the two flagella of the free-living proto-apicomplexans. We believe that future analysis of the localization of the homologs of components of additional structural landmarks, such as the apical polar ring (the organizing center for the cortical MT array), will be highly informative on the evolution of the cytoskeletal architecture of the apicomplexans.

The successful transformation of *Chromera* now paves the way for developing additional genome editing tools such as homologous recombination or CRISPR-Cas9. The prospect is exciting. For instance, the analysis of function, in free-living *Chromera,* of homologs of genes known to be critical for apicomplexans’ parasitism will reveal the ancestral cellular activities from which behaviors essential to the intracellular parasitic lifestyle arose, such as the ability to invade into another cell using gliding motility. Parallel structural and function analyses of the conoid in *Toxoplasma* and pseudoconoid in *Chromera* will provide insights into the evolution of cellular structures (e.g., What kind of molecular inventions allow *Toxoplasma* but not chromerids to override the intrinsic property of the tubulin subunits to form a tube?). It will also help to understand how specific structural transitions (e.g., pseudoconoid to conoid) are related to the transition from a free-living to an intracellular parasitic lifestyle. We believe our work marks a major step of developing *Chromera* into a tractable model system to finally answer these questions.

## MATERIALS AND METHODS

### Chromera velia cultures

*Chromera velia* (CCMP2878) was obtained from the National Center for Marine Algae and Microbiota at Bigelow Laboratory for Ocean Sciences. The growth media of the culture is L1 media without silicon prepared with the L1 media kit (MKL150L, Bigelow Laboratory for Ocean Sciences) [59]. The cultures were maintained in T25 or T75 flasks in an environmental chamber (1-35VL, Percival Scientific, IA, USA) with a light-dark cycle of 14 hr light:10 hr dark.

*Plasmids construction* (Sequences for all synthesized DNA fragments are included in Supplemental File 1) The pCvGAPDH and pCvL31 plasmids were based on plasmid backbones constructed by Genscript (Piscataway, NJ, USA). For generating these plasmid backbones, DNA fragments containing the putative promoter region, kozak sequence, CDS for mNeonGreen, and 3’UTR were synthesized and then ligated into the BglII and AflII sites of a pBluescript (PBS). Full sequences of pCvGAPDH-mNeonGreen, pCvL31-mNeonGreen are included in Supplemental File 1. The kozak sequence for CvGAPDH (gctcagaagATG was used for all constructs described below.

*pCvGAPDH-NanoLuc*: The CDS for *NanoLuc* was amplified from pCDH-EF1-Nluc (#73024, Addgene) using the primers P1 and P2 (Supplemental Table 1) and inserted between the *Bgl*II and *Afl*II sites of *pCvGAPDH-mNeonGreen* to replace the CDS of mNeonGreen.

*pCvGAPDH-NanoLuc*^opt^: A gBlock DNA fragment that includes the CDS for NanoLuc codon optimized for *Chromera velia* with an intron between the promoter and the kozak sequence was synthesized (IDT Inc.) and inserted between the *Afl*II and *Nsi*I sites of *pCvGAPDH-NanoLuc*. The intron introduced is the first intron of a *Chromera* β-tubulin (VEupathDB: *Cvel_33153*). For codon-optimization, the codon usage table for *Chromera* was obtained from the High-performance Integrated Virtual Environment-Codon Usage Tables (HIVE-CUTs) database [60, 61], which was then imported into SnapGene to replace the non-preferred codons based on the *Chromera* codon usage table.

*pCvL31-NanoLuc*^opt^: The sequence used as promoter was amplified from *pCvL31-mNeonGreen* using the primers P3 and P4, and the DNA fragment that includes the CDS for NanoLuc codon optimized for *Chromera velia* with an intron between the promoter and kozak sequence was amplified using the primers P5 and P6. These two fragments were fused using overlap extension PCR using the primers P4 and P5. The fused fragment was inserted between the *Pst*I and *AflI*I sites of *pCvL31-mNeonGreen* using Hi-Fi assembly.

*pCvGAPDH-CvH2B-mStayGold*: A gBlock fragment that includes the kozak sequence, CDS of Cvhistone H2B(Cvel_19200), a linker (Supplemental Table 1), and mStayGold codon-optimized for *Chromera*, with both ends containing ∼20-bp sequence that overlaps with the backbone insertion sites was synthesized (IDT Inc.) and inserted into the *Pst*I and *Afl*II sites of *pCvGAPDH-NanoLuc*^opt^ using Hi-Fi assembly.

*pCvGAPDH-CvACP-mStayGold, pCvGAPDH -CvIMC13-mStayGold*: For all these constructs, a PstI-5’ and *Bgl*II-3’ flanked DNA fragment that includes the kozak sequence, CDS of the target gene, and a linker (Supplemental Table 1) was synthesized and ligated into the *Pst*I and *Bgl*II sites of the *pCvGAPDH-CvH2B-mStayGold* plasmid. The VEupathDB accession numbers of these genes are as follows: *CvACP (Cvel_32060), CvIMC13 (Cvel_9514).* Synthesis of DNA fragments and cloning were carried out by Genscript (Piscataway, NJ, USA)

*pCvGAPDH-mStayGold-CvTubA1*: A *Pst*I-5’ and *Afl*II-3’ flanked DNA fragment that includes the kozak sequence, CDS of mStayGold, a linker (Supplemental Table 1) , CDS of *CvTubA1* (Cvel_22324) with stop codon was synthesized and ligated into the *Pst*I and *Afl*II-sites of the *pCvGAPDH-CvH2B-mStayGold* plasmid. Synthesis of the DNA fragment and cloning were carried out by Genscript (Piscataway, NJ, USA)

### Toxoplasma cultures and transfections

*Toxoplasma* tachyzoite cultures were maintained in Human Foreskin Fibroblasts (HFFs) as previously described [28–31]. *Toxoplasma* transfections were carried out using a BTX ECM 630 (Harvard Apparatus) and 2mm gap cuvettes based on established protocols [28–31]. Transfected parasites were inoculated in MatTek glass-bottom dishes (MatTek Corporation, CAT# P35G-1.5-14-C or CAT# P35G-1.5-21-C) with rat aorta cells (A7r5 ; ATCC# CRL-1444) cells and imaged ∼ 24 to 36 hours after transfection.

### Chromera transfections

#### Cell preparation

Freshly saturated *Chromera* cultures (OD600 ≈ 1.5–2.0) were diluted 1:2 or 1:3 in fresh L1-Si medium and incubated for 4-6 days, until reaching an OD600 of 0.8–1.3. Only cultures aged 6 days or less were used for optimal transfection efficiency. Cell concentration was determined using a hemocytometer, and 2.5 × 10^6^ cells were prepared for each transfection. Cells were pelleted by centrifugation at 3000 × g for 3-5 min and washed four times in 100 µL of room temperature MAX Efficiency™ Transformation Reagent for Algae (“MEB”, A24229, Invitrogen) for every 1 mL of culture harvested. After the final wash, cells were resuspended in 50 µL of transformation reagent per 2.5 × 10^6^ cells per transfection.

#### DNA preparation

The expression plasmid was linearized using *NotI*-HF, *Sca*I-HF, or double-digested with *Hind*III-HF and *Spe*I-HF (NEB) and complete digestion was confirmed by DNA electrophoresis. Double-digestion cuts the plasmid into two parts: the expression cassette, and the cloning backbone. This method produced a smaller cassette that may aid delivery and expression. The linearized plasmid was precipitated by adding 0.1 volume of 3 M sodium acetate and 3 volumes of ice-cold 100% ethanol before incubation at -20°C for at least 30 min. The DNA was pelleted by centrifugation at ∼21,000 × g for 30 min, then washed once with 70% ethanol and centrifuged at the same speed for 10 min. The pellet was air-dried before resuspending with nuclease-free water at a target concentration of at least 4 mg/mL. The DNA was prepared at a high concentration to minimize the amount of water added to the transfection mix.

#### Electroporation

Electroporation was mainly carried out with the NEPA21 (Bulldog Bio) electroporator, which generates square wave poring and transfer pulses. Poring pulses deliver higher energy and are designed to create pores for plasmids to cross the plasma membrane. Transfer pulses, at lower energy, are designed to transport and retain the DNA molecules in the cell after the poring pulses. The voltage, length, number, interval, decay rate and polarity switching of the pulses can all be set by the user. As *Chromera* coccoids have a thick cell wall that likely impede DNA delivery, instead of the 2mm gap cuvettes typically used for eukaryote transfections. we used 1mm gap cuvettes in all transfections to increase strength of the electric field. Transfer pulse with the following parameters were kept consistent throughout the transfection trials: 25 V, pulse length of 50 ms, pulse interval of 50 ms, 10 pulses with 40% decay rate and polarity switch.

The plasmid (and carrier DNA with stock concentration of at least 10 mg/mL, if using) was added to the cell suspension and mixed gently before pipetting into the electroporation cuvette. Immediately after the pulse was delivered, 100 µL of recovery media (0.5 M sorbitol in L1-Si) was added to the cuvette, and the whole suspension was transferred to a 24-well plate (for NanoLuc constructs) or 35 mm glass-bottom MatTek dish (for fluorescence constructs) containing 700 µL recovery media. The plates or dishes were incubated in the dark for 24 h before incubation under normal conditions.

To detect NanoLuc activity, 1/8^th^ of the total transfected cells was pelleted and resuspended in 50 µL of assay buffer (0.06% NP-40, 0.2% gelatin, 4 mM thio-urea, 0.8 mM DTT in 100 mM Tris-Cl pH 8.0) and vortexed for 5 sec. The tubes were immersed in a sonicator bath for 5-10 sec, then pipetted into Corning® 96-well Solid White Flat bottom plates. Immediately before luminescence reading, 50 µL of 0.1 µM furimazine in assay buffer was then added and the plate was gently shaken. The plate was inserted into the BioTek Synergy H1 reader and luminescence was read after a 30 sec delay with 2 s integration time and 3-6 mm probe height. Each time after a sample was taken out for measurement, fresh L1-Si was added to the original culture to replenish the media.

Conditions that consistently produced transfected cells expressing fluorescent proteins are as follows: For each transfection, 2.5 × 10^6^ cells electroporated with 16 µg of plasmid double-cut with *Hind*III and *Spe*I, without carrier DNA, electroporated using the following poring pulse parameters: poring voltage 300 V, pulse length of 10 ms, pulse interval of 50 ms, 2 pulses with 20% decay rate and polarity switch, and transfer pulses as described above. Transfected cells were typically imaged 4 days or more after transfection.

### Fluorescence imaging of live cells

For *Chromera* imaging, following transfection and 24 h of incubation in the dark, 2 mL of L1-Si media was added to the dishes containing transfected cells, Cells in dishes were transferred into the environmental chamber and grown under normal conditions. After 3 days (4^th^ day post-transfection), cells were screened, and images were taken using an OMX Flex microscope with 60X oil objective lens (Olympus PlanApo N 60x/1.42 Oil). *Toxoplasma* imaging was carried out on DeltaVision and OMXflex imaging systems as described before [31, 52, 62]. Contrast levels were adjusted to optimize the display.

### Expansion microscopy (ExM)

ExM gels containing *Toxoplasma* were prepared as described in [52, 62]. For *Toxoplasma* samples, tubulin was labeled with a mouse monoclonal anti-acetylated tubulin (T6793-6-11B-1, Sigma-Aldrich, 1:250) followed by goat anti-mouse IgG 568 (A11031, Molecular Probes, 1:400). MORN1 was labeled using a rat anti-TgMORN1 antibody (1:200) [46] and followed by goat anti-rat IgG 488 (11006, Molecular Probes, 1:400).

For *Chromera* ExM samples, *Chromera* cells were harvested at peak flagellation time during the light phase of the light-dark cycle [22] by centrifugation at 3000 x g for 3 min and the pellet was resuspended in 50-80 mM of sodium acetate in L1-Si media to immobilize the flagellates. Onto a gelatin-coated 12mm round coverslips, ∼250 µL of the concentrated cells were pipetted, and incubated at room temperature in a humid chamber for 10-30 min. The remaining cells in solution were removed carefully before fixing the coverslip in 2% formaldehyde in L1-Si for 5 minutes. Excess solution was blotted and the coverslip washed with PBS gently before transferring into the ExM anchoring solution [52, 62] followed by incubation for 3h at 37°C in a humid chamber. The rest of ExM was performed as previously described [52, 62].

The *Chromera* ExM samples were labeled with a cocktail of mouse monoclonal anti-tubulin antibodies consisting of anti-acetylated tubulin (T6793-6-11B-1, Sigma-Aldrich, 1:150), anti-beta tubulin (T5293, Sigma-Aldrich, 1:200), and anti-alpha tubulin (T6074, Sigma-Aldrich, 1:150), followed by a goat anti-mouse IgG AF488 (A11029, Invitrogen, 1:350) and a rat anti-TgMORN1 antibody (1:150) [46] followed by goat anti-rat IgG Cy3 (112-165-167, Jackson ImmunoResearch Labs, 1:350).

Fully expanded gels were imaged using an OMX Flex imaging system with 60X oil objective lens (Olympus PlanApo N 60x/1.42 Oil) as described in [52, 62]. Contrast levels were adjusted to optimize the display.

## Acknowledgments

We are grateful to Dr. Bradley Olson (Augusta University) for the suggestion of including an intron into the *Chromera* expression plasmid. We thank Drs. John M. Murray, Luisa F. Arias Padilla, and Jonathan Munera Lopez for helpful discussions and Matea Susac for tissue culture support.

