## Supplemental File 1 for "Transfection of the free-living alga *Chromera velia* enables direct comparisons with its parasitic apicomplexan relative, *Toxoplasma gondii*"

>pCvL31-mNeonGreen (5292 bp)
CACCTAAATTGTAAGCGTTAATATTTTGTTAAAATTCGCGTTAAATTTTTGTTAAATCAGCTCATTTTTTAACCAATAGG
CCGAAATCGGCAAAATCCCTTATAAATCAAAAGAATAGACCGAGATAGGGTTGAGTGTTGTTCCAGTTTGGAACAAGAGT
CCACTATTAAAGAACGTGGACTCCAACGTCAAAGGGCGAAAAACCGTCTATCAGGGCGATGGCCCACTACGTGAACCATC
ACCCTAATCAAGTTTTTTGGGGTCGAGGTGCCGTAAAGCACTAAATCGGAACCCTAAAGGGAGCCCCCGATTTAGAGCTT
GACGGGGAAAGCCGGCGAACGTGGCGAGAAAGGAAGGGAAGAAAGCGAAAGGAGCGGGCGCTAGGGCGCTGGCAAGTGTA
GCGGTCACGCTGCGCGTAACCACCACACCCGCCGCGCTTAATGCGCCGCTACAGGGCGCGTCCCATTCGCCATTCAGGCT
GCGCAACTGTTGGGAAGGGCGATCGGTGCGGGCCTCTTCGCTATTACGCCAGCTGGCGAAAGGGGGATGTGCTGCAAGGC
GATTAAGTTGGGTAACGCCAGGGTTTTCCCAGTCACGACGTTGTAAAACGACGGCCAGTGAATTGTAATACGACTCACTA
TAGGGCGAATTGGAGCTCCACCGCGGTGGCGGCCGCTCTAGAACTAGTGGATCccatcgactcaggagaggggcacagaa
ggaatgaagagtcaggaggtggggcagaagttaagaaaaggggggaatgtgaggaatggtttgaggggggttaagagtag
attatgtaagagagagaagttattcctcactgaggtcggatcagcagttatagccagacattggcctgaaaaagaggtgg
agggaagaagggaagcagaagaagtgggttaagtgacgacagcagaagggaaaaacacctaggatgatcatcacacatga
agccagcacacaaacaaagagagtctgcctcccttcctgtccgcccgagggagagaagagttcctccaggcttcactcaa
gcatttgcagacttccaatcgctcgtccgtgcaggctttcctagtacccccaaacaaaaggtaccgagttccatcaatgg
tcgcttaattctgcccctccgtcagctcagtctgtcgtgaggggactgttcagttcaacgcccagccagacgccttaagt
tatgacccaccggaaccagttccaccagacccggtacctcccttgtacagctcgtccatgcccatcacatcggtaaaggc
cttttgccactccttgaagttgagctcggtcttggagtgcttgagctccgtcttacggaacacgtacatcggctggttct
tcagatagttagccgccattggcttggcaaaggtgtaggtggtccgcgcagtgctccggtagcgcttgccatttccagtg
gtgtaactccacttaaaggtactgatgatggttttgtcgttggggtaagtcttcttcgacctgcaccagtccgcagcggt
cagcgagttggtcatcacaggaccgtcagcagggaaaccagtccccttcacctgggcctctcctttgatgtggcttccct
cgtaggtgtagcggtagttaacagtaagggaggcaccatcttcaaactgcattgtgcgatggacttggtagccggagcca
tctaccatggcggcctggaaaggcgacatcccgtcagggtagggcaggtactgatggaagccatacccgatatgagggac
cagaatccaggggggagaactggaggtcacccttggtggacttcaggtttaactcctcataaccatcatttggattgccgg
tgccctgacccaccatgtcaaagtccacaccgttgatggagccaaagatgtgtaactcatgtgtcgctgggagagaggcc
atgttatcctcctcgcccttgctcaccatctttctcggtccctctttgatccgaatgtgcgcaggagcgaaggcgaagaa
gctcttcaaatcacacaaacggatcgaggcgtttccgtcgatccgattgtgtggagaaatttggctccaccccggatgat
tttctcgtggcacccctggcgacctttcggccgcgaaaattcatctcgcggctctgtactgcgcggggacgaaggggcac
aaaaaactttccgggcccactcggaattgtctttccgggacatcatgcacccaaaatcttctccgtctcgtagacctggt
attatatataccgctctgattttgaattcctttgtgtgagaggctgcagcactcttacgcaaatactgaataccgtcgct
ttcttgagtgtcgatctgcttgctggcggcttcccagcttctgttttgccataagggtccctgcaatgaagtcaatcctc
tgcgtggctgctcttgccgtcacagggactctccctgtctccgaggccttcacccccctcttggccaccaaagcgccttc
cagggcagaggcgacttctcgcaaggcagtggcagacatggtgggggttgatacggattctgggaacacagtttgggacc
ctcttgacattgcatccaagtgcgagtttaatcgcaagaaggggcgggacaacgtgatttttttcaacctggcctcatctg
aactggatgagagaggcagagctgaagcacggacgcatctgtatgcttgcggttctgggtgttctgttcacatagcttgg
catcaaaattgggggtttcatttcaaattcctctactgactggtgggagaacctcggccgcgtgtgggctgagcatcctg
aagtgattgccttcatcatcatggcgggagggttcattgacatggccacttacgacactcgcatgtggtcaggcaacaac
aagcgcgaggctgggaacctgggctttgatccgttgaacttgatggcgaaggacgacaatttcgccctcaaggagctcaa
gcacggtgggtgtttttgctgtctgtttcatgcatgcgcatggttggtctggtactcctcgatgttcctttccgtttgaa
tcggctgctcggAAGCTTATCGATACCGTCGACCTCGAGGGGGGGCCCGGTACCCAGCTTTTGTTCCCTTTAGTGAGGGT
TAATTTCGAGCTTGGCGTAATCATGGTCATAGCTGTTTCCTGTGTGAAATTGTTATCCGCTCACAATTCCACACAACATA
CGAGCCGGAAGCATAAAGTGTAAAGCCTGGGGTGCCTAATGAGTGAGCTAACTCACATTAATTGCGTTGCGCTCACTGCC
CGCTTTCCAGTCGGGAAACCTGTCGTGCCAGCTGCATTAATGAATCGGCCAACGCGCGGGGAGAGGCGGTTTGCGTATTG
GGCGCTCTTCCGCTTCCTCGCTCACTGACTCGCTGCGCTCGGTCGTTCGGCTGCGGCGAGCGGTATCAGCTCACTCAAAG
GCGGTAATACGGTTATCCACAGAATCAGGGGATAACGCAGGAAAGAACATGTGAGCAAAAGGCCAGCAAAAGGCCAGGAA
CCGTAAAAAGGCCGCGTTGCTGGCGTTTTTCCATAGGCTCCGCCCCCCTGACGAGCATCACAAAAATCGACGCTCAAGTC
AGAGGTGGCGAAACCCGACAGGACTATAAAGATACCAGGCGTTTCCCCCTGGAAGCTCCCTCGTGCGCTCTCCTGTTCCG
ACCCTGCCGCTTACCGGATACCTGTCCGCCTTTCTCCCTTCGGGAAGCGTGGCGCTTTCTCATAGCTCACGCTGTAGGTA
TCTCAGTTCGGTGTAGGTCGTTCGCTCCAAGCTGGGCTGTGTGCACGAACCCCCCGTTCAGCCCGACCGCTGCGCCTTAT
CCGGTAACTATCGTCTTGAGTCCAACCCGGTAAGACACGACTTATCGCCACTGGCAGCAGCCACTGGTAACAGGATTAGC
AGAGCGAGGTATGTAGGCGGTGCTACAGAGTTCTTGAAGTGGTGGCCTAACTACGGCTACACTAGAAGGACAGTATTTGG
TATCTGCGCTCTGCTGAAGCCAGTTACCTTCGGAAAAAGAGTTGGTAGCTCTTGATCCGGCAAACAAACCACCGCTGGTA
GCGGTGGTTTTTTTGTTTGCAAGCAGCAGATTACGCGCAGAAAAAAAGGATCTCAAGAAGATCCTTTGATCTTTTCTACG
GGGTCTGACGCTCAGTGGAACGAAAACTCACGTTAAGGGATTTTGGTCATGAGATTATCAAAAAGGATCTTCACCTAGAT
CCTTTTAAATTAAAAATGAAGTTTTAAATCAATCTAAAGTATATATGAGTAAACTTGGTCTGACAGTTACCAATGCTTAA
TCAGTGAGGCACCTATCTCAGCGATCTGTCTATTTCGTTCATCCATAGTTGCCTGACTCCCCGTCGTGTAGATAACTACG
ATACGGGAGGGCTTACCATCTGGCCCCAGTGCTGCAATGATACCGCGAGACCCACGCTCACCGGCTCCAGATTTATCAGC
AATAAACCAGCCAGCCGGAAGGGCCGAGCGCAGAAGTGGTCCTGCAACTTTATCCGCCTCCATCCAGTCTATTAATTGTT
GCCGGGAAGCTAGAGTAAGTAGTTCGCCAGTTAATAGTTTGCGCAACGTTGTTGCCATTGCTACAGGCATCGTGGTGTCA
CGCTCGTCGTTTGGTATGGCTTCATTCAGCTCCGGTTCCCAACGATCAAGGCGAGTTACATGATCCCCCATGTTGTGCAA
AAAAGCGGTTAGCTCCTTCGGTCCTCCGATCGTTGTCAGAAGTAAGTTGGCCGCAGTGTTATCACTCATGGTTATGGCAG
CACTGCATAATTCTCTTACTGTCATGCCATCCGTAAGATGCTTTTCTGTGACTGGTGAGTACTCAACCAAGTCATTCTGA
GAATAGTGTATGCGGCGACCGAGTTGCTCTTGCCCGGCGTCAATACGGGATAATACCGCGCCACATAGCAGAACTTTAAA
AGTGCTCATCATTGGAAAACGTTCTTCGGGGCGAAAACTCTCAAGGATCTTACCGCTGTTGAGATCCAGTTCGATGTAAC
CCACTCGTGCACCCAACTGATCTTCAGCATCTTTTACTTTCACCAGCGTTTCTGGGTGAGCAAAAACAGGAAGGCAAAAT
GCCGCAAAAAAGGGAATAAGGGCGACACGGAAATGTTGAATACTCATACTCTTCCTTTTTCAATATTATTGAAGCATTTA
TCAGGGTTATTGTCTCATGAGCGGATACATATTTGAATGTATTTAGAAAAATAAACAAATAGGGGTTCCGCGCACATTTC
CCCGAAAAGTGC

>pCvGAPDH-mNeonGreen (5432 bp)
cacctaaattgtaagcgttaatattttgttaaaattcgcgttaaatttttgttaaatcagctcattttttaaccaatagg
ccgaaatcggcaaaatcccttataaatcaaaagaatagaccgagatagggttgagtgttgttccagtttggaacaagagt
ccactattaaagaacgtggactccaacgtcaaagggcgaaaaaccgtctatcagggcgatggcccactacgtgaaccatc
accctaatcaagttttttggggtcgaggtgccgtaaagcactaaatcggaaccctaaagggagcccccgatttagagctt
gacggggaaagccggcgaacgtggcgagaaaggaagggaagaaagcgaaaggagcgggcgctagggcgctggcaagtgta
gcggtcacgctgcgcgtaaccaccacacccgccgcgcttaatgcgccgctacagggcgcgtcccattcgccattcaggct

gcgcaactgttgggaagggcgatcggtgcgggcctcttcgctattacgccagctggcgaaagggggatgtgctgcaaggc
gattaagttgggtaacgccagggttttcccagtcacgacgttgtaaaacgacggccagtgagcgcgcgtaatacgactca
ctatagggcgaattggagctccaccgcggtggcggccgctctagaactagtgGATCcaaagctagaaacaTagcttcaat
gaaagtttcaatgatttagaaaagttttagcaaacttcattgtcgctacaagccagtcaagtctttgaatgccgaaaca
ttcggtcaacctcaaagctacttcagacttcttctaatagtttcactttcaagggtgagccgtcagtcagagacttgaaa
aaactcacgcgtcccaactcgactgagcgacagaagaacagacaaaaccgctcaaacacactcgacaaagtaatgtttaa
ggcttactactaaaccgggaagaccgcggcgggaagatcgcacacagtcacgcggtggaaagaaaaatcagccaagcgac
tgactgaaaaggtagaagggggaatcaatacgaatggaaatcgcgacaccacctctcttctctcaccagcaatcgaggcc
cacttcccttcctttcggtttcaaaacaaagagaagagagagaccagactaagacagacaaaacagacagagaggtgaga
caaagaactgcgatcgtttgaaaaacgttaattgaaagcatctgcaacgcattgagcatcatgcactcaacaaacaactg
cgttaacagccagccggccggcggttgccaggcaaacaagcaagctgcCTTAAGttatgacccaccggaaccagttccac
cagacccggtacctcccttgtacagctcgtccatgcccatcacatcggtaaaggccttttgccactccttgaagttgagc
tcggtcttggagtgcttgagctccgtcttacggaacacgtacatcggctggttcttcagatagttagccgccattggctt
ggcaaaggtgtaggtggtccgcgcagtgctccggtagcgcttgccatttccagtggtgtaactccacttaaaggtactga
tgatggttttgtcgttggggtaagtcttcttcgacctgcaccagtccgcagcggtcagcgagttggtcatcacaggaccg
tcagcagggaaaccagtccccttcacctgggcctctcctttgatgtggcttccctcgtaggtgtagcggtagttaacagt
aagggaggcaccatcttcaaactgcattgtgcgatggacttggtagccggagccatctaccatggcggcctggaaaggcg
acatcccgtcagggtagggcaggtactgatggaagccatacccgatatgagggaccagaatccagggggagaactggagg
tcacccttggtggacttcaggtttaactcctcataaccatcatttggattgccggtgccctgacccaccatgtcaaagtc
cacaccgttgatggagccaaagatgtgtaactcatgtgtcgctgggagagaggccatgttatcctcctcgcccttgctca
cAGATCTcttcatcttctgagcgagacggagggaaaaggcgtggaaggaacgacaccgtgtgctgtactaccgtgactct
gtctttggacccgacggattggcaggaaaggagctgggacacaagagaaaaaagagaggttgaaaatggcaaagagacgg
tgtctccaaaaattgcgcggcccgcgcgaaacgtgaccgttatttgtgcatgtgcatgcatgggcgtggcccccgcctct
ttcggcgcctctggtccttgaaaagtgcggaaattcaaggccacggcatgccttgttagaggttgagttccttggatgat
tctccagggaatattgggccctggcagccggccccagaccgcagatttgcggctacgtgtacagtcgaccatgcacgcgc
gcggtcccgaaaggcggacacgcggggatgtgaatcatggagacggcaatagaattacgagattccatgagttccgtcta
tcgaatcgcaagattcggccgatcagtaactgtctctattccaacgatgccttgacagcacatcacgagagcaaacctgc
ctgcacgcgatctgacttgccacgtgggggcgcagcctattaatctttcaagtagtcgttcatcactccgcAcgagcatt
tctttctgatatcctcaccgtggtgcgtgtgttgcttgagcacatccgctcgccagctctgtgagcagagaccacagaaa
aaaggaaaatggacccacgatttccaggcaacagccagcagggtggatttggcacgggaggtagcggtccgggtggtttt
gactcaaacaatcccccctccgatcgatccccagtctcaggctacggggaaccctgaaagccagccgcttgtcgagaatgg
cgagaggatcgaaaacatcctcgcaggagcgaactgcacccacaaaattctgtttttcgtctgttccgtacttttgctct
gctgcgcgattgtggggcttttgacatcgcttgcgacgtttgagattggggagcttgtcgatgacatctttctcatcacc
ttcactgtcctcctcatttttctcaacttccccaaagccaccgttgaccacgcggtcaaggcatcgaaatggAagcttat
cgataccgtcgacctcgagggggggcccggtacccagcttttgttccctttagtgagggttaattgcgcgcttggcgtaa
tcatggtcatagctgtttcctgtgtgaaattgttatccgctcacaattccacacaacatacgagccggaagcataaagtg
taaagcctggggtgcctaatgagtgagctaactcacattaattgcgttgcgctcactgcccgctttccagtcgggaaacc
tgtcgtgccagctgcattaatgaatcggccaacgcgcggggagaggcggtttgcgtattgggcgctcttccgcttcctcg
ctcactgactcgctgcgctcggtcgttcggctgcggcgagcggtatcagctcactcaaaggcggtaatacggttatccac
agaatcaggggataacgcaggaaagaacatgtgagcaaaaggccagcaaaaggccaggaaccgtaaaaaggccgcgttgc
tggcgtttttccataggctccgcccccctgacgagcatcacaaaaatcgacgctcaagtcagaggtggcgaaacccgaca
ggactataaagataccaggcgtttccccctggaagctccctcgtgcgctctcctgttccgaccctgccgcttaccggata
cctgtccgcctttctcccttcgggaagcgtggcgctttctcatagctcacgctgtaggtatctcagttcggtgtaggtcg
ttcgctccaagctgggctgtgtgcacgaaccccccgttcagcccgaccgctgcgccttatccggtaactatcgtcttgag
tccaacccggtaagacacgacttatcgccactggcagcagccactggtaacaggattagcagagcgaggtatgtaggcgg
tgctacagagttcttgaagtggtggcctaactacggctacactagaaggacagtatttggtatctgcgctctgctgaagc
cagttaccttcggaaaaagagttggtagctcttgatccggcaaacaaaccaccgctggtagcggtggtttttttgtttgc
aagcagcagattacgcgcagaaaaaaaggatctcaagaagatcctttgatcttttctacggggtctgacgctcagtggaa
cgaaaactcacgttaagggattttggtcatgagattatcaaaaaggatcttcacctagatccttttaaattaaaaatgaa
gttttaaatcaatctaaagtatatatgagtaaacttggtctgacagttaccaatgcttaatcagtgaggcacctatctca
gcgatctgtctatttcgttcatccatagttgcctgactccccgtcgtgtagataactacgatacgggagggcttaccatc
tggccccagtgctgcaatgataccgcgagacccacgctcaccggctccagatttatcagcaataaaccagccagccggaa
gggccgagcgcagaagtggtcctgcaactttatccgcctccatccagtctattaattgttgccgggaagctagagtaagt
agttcgccagttaatagtttgcgcaacgttgttgccattgctacaggcatcgtggtgtcacgctcgtcgtttggtatggc
ttcattcagctccggttcccaacgatcaaggcgagttacatgatcccccatgttgtgcaaaaaagcggttagctccttcg
gtcctccgatcgttgtcagaagtaagttggccgcagtgttatcactcatggttatggcagcactgcataattctcttact
gtcatgccatccgtaagatgcttttctgtgactggtgagtactcaaccaagtcattctgagaatagtgtatgcggcgacc
gagttgctcttgcccggcgtcaatacgggataataccgcgccacatagcagaactttaaaagtgctcatcattggaaaac
gttcttcggggcgaaaactctcaaggatcttaccgctgttgagatccagttcgatgtaacccactcgtgcacccaactga
tcttcagcatcttttactttcaccagcgtttctgggtgagcaaaaacaggaaggcaaaatgccgcaaaaaagggaataag
ggcgacacggaaatgttgaatactcatactcttcctttttcaatattattgaagcatttatcagggttattgtctcatga
gcggatacatatttgaatgtatttagaaaaataaacaaataggggttccgcgcacatttccccgaaaagtgc

>gblock (pCvGAPDH–CvH2B–mStayGold)  (1120 bp)
aaacaagcaagctgcCTTAAGTTAGAGGTGGGCCTCGAGGGTCTCGGACTGGATGATGTGGTCTCTCTCCTCGGTGTCGT
CCTTGGACTGGGTGTACTGCTTTCTGATCCAGTGGAAGGGCACGTCGGGGGCGGGCTGGTTGTGGAGGGGCTTGATGATG
GTGTTCTGGTAGGCCTCCACGCACTTGGACTCGTCGGAGAGGAGGGGGTACTGGAGGGTCACGGTGCACTCCACCCCGTC
GTCGATGGGGATGTGCTGCACCTCGTTGGGGAGGGACACGTCCATGTCCCCGGTGAGCACGGGGGAGTTCTCCTTGAAGT
CCTGCCCGGTGAACTCCACGATGTTGTGGTAGGTCCCGTTCTTCATGAAGTGCTGGTGCTTGGCGTGGATGGACCCGTCC
CCCTTGTACTGGATGTGTCTGTCGTAGGTGAACCCCTCGGGCATCACCTCGTGGAACCAGTTCTTGAGCCCGGAGGGGTA
CTTGGTGTAGTACTTCATCCCGTACCCGAAGGAGGTCCCGAGGGCGGCCCAGGACATGGGGAGCTTCCCGGAGGTGCACA
CGTACTTCCCCTTGTGGGACCCCTCGTGGGAGTTCCCCTCCCCCTCCCCCTCCACGGTGAAGGACTTCCCGTTGATGGTC
CCCTTGAGCTGGAACTTGAAGGGCACCACCCCGGTGAAGAGCTCCTCCCCGGTGGACACagatctacctgaaccagttcc
accagacccggtcccgggCGAAGAGAACTTCGTGACAGCCTTGGTTCCCTCAGAGATGGCGTGCTTGGCGAGTTCACCAG
GCAGGACCAGGCGGATGGCTGTCTGGACCTCTCTGCTGGAGAGCGTGTCCTTCTTGTTGATCTTGCAGAGCTTTCCTGCC

TCCACAGCGATCTTCTCGAAGGTGTCGTTGATGAACGAGTTCATGATCGACATGCTCTTCTTGGAGATTCCAGTCTCAGG
GTGGACCTGCTTCAACACCTTGTAGATGTAGGAAGCGAAGGACTCAACCCTCTTGGACTTGCGCTTCTTGCCGCCCTTGG
GCACGGTCGCCTTCTTGGCGGACTTCTTGGATGCCTTAGCTTTGGGTGCCATcttctgagcctgcagagccgagggagaa

>gblock (pCvGAPDH–NanoLuc[opt])  (863 bp)
aacaagcaagctgcCTTAAGTTAGGCGAGGATTCTCTCGCAGAGTCTccaCCCggtcacCCCgttgatggtCACTCTgaa
GAGGAGggaCCCgtcggggttgatGAGTCTctcgtcgatGATCTTgttCCCgttccaGAGggtcccGGTCACGGTgatct
tCTTCCCgtcgaacacggcgatCCCCTCGTAGGGTCTCCCgaaGTAgtcgatcatgttGGGGGTCACcccgtcgatCACG
AGGGTCCCGTAgtgGAGgatcaccttGAAgtgGTGGTCGTCcacGGGgtacaccaccttGAAGATCTTCTCgatctgCCC
catCTGgtcCCCGGAGAGCCCCTCGTAGGGgatgatCACGTGgatgtcgatcttGAGcccGTTCTCCCCGGAGAGCACGA
TTCTCTGgatGGGGGTCACggacaccccgagGTTctgGAAGAGGGAggacacCCCCCCctgCTCGAGCACCTGgtcGAGg
ttgtaCCCggcGGTctgTCTccagtccccCACgaaGTCCTCgagGGTgaaCACcatcttctgagcctgcagagccgaggg
agaagaaaagaaaatttcagcaaaggaaacgtgatcaacacagccagaacaaaaaatggaaagtaggagagagtttgagg
agaagactgacgagacggagggaaaaggcgtggaaggaacgacaccgtgtgctgtactaccgtgactctgtctttggacc
cgacggattggcaggaaaggagctgggacacaagagaaaaaagagaggttgaaaatggcaaagagacggtgtctccaaaa
attgcgcggcccgcgcgaaacgtgaccgttatttgtgcatgtgcatgcatgggcgtggccccc

>gblock (pCvGAPDH–CvACP–mStayGold)  (429 bp)
ctgcaggctcagaagATGATGAGCCGTCTTCTTTGCGCTGCCGCTCTTCTCCTTTGTGGAGCATCTGCATTCGACCTGGCATTTGTCTCTCCCCC
CTCCGCTCTCAGGGGCTTTCGGCGCGCACCCCAGGTGACAATTCTGCGCATGGCCGACTCTGACGATGAAATCTTGGACAAGGTGCGCGCAGTGG
TTGCCGACCAGCTGGGCGTTGCTGCCGAGAAGGTTGTGATGGAGGCTGCCTTTGTGGAGGACTTGGGGGCTGACTCTCTGGACTCTGTTGAGCTC
GTGATGGCATTGGAGGAGAAGTTCGGAATTGAGATTCCCGATGAGAGCGCCACCAAGATaTCCACTGTCAAGGCTGCTGTGGACTACATCAAGGC
CAACCAGAAGcccgggaccgggtctggtggaactggttcaggtagatct

>gblock (pCvGAPDH–CvIMC13–mStayGold)(1326 bp)
ctgcaggctcagaagATGTCAGGTCCAGGCCTCGTCACCAATGCATACGGCAAGGATATGTCTCCTGCCCCTGTCCCGCAGCAGACCGGTGCCAT
GCCGGTTGGTTACCCCCCCCAGCCAGGAATGGTTCCTGGGTACGTGCCTCAGTACGGGGAAGCTCGGACCGTTGGATACGGCCAGGGCTCTACTG
TTGGAATGGCCCATGCGGGATTTGTCATGACAGTTGAGAAGGTTGTTGAGGTCCCCCAGGTGCAGATTGTCGACAAGGTCGTGGAGGTGCCCGAA
ATCCAGGAGGTCGTCAAGCACGTTCCCCGCCTGGAGGTTCAGGAGGTCACTCGGGAAGTCCACGTCCCAGAGCTGGAGGTGGTGGAGAAGACGGT
CGAGGTGCCCCACGTTCAGGTCGTTGACAAGTACGTTGAGGTCCCGCAGGTACAGGAGGTCATCAAACACGTCCCCAAAATCATCCAGCAGGAAG
TGCTCAGGGAGTACGTGCGGCATGTGCCGAAAGTAGAGGTACGCACAGTGGAGGTCATCAAGCAGGTGCCCGTGCCACAGTACGTAGATCAGCCC
TACGAGGTCATCAAACATGTACCCAAACCCGTCCCCGTGTTCCACCCTCGCCATGTCCCTGTGCCGAAGGTCGTCACCGTCGACAGGCCCTACCA
GGTTCATGTTCCTCAGATCGTCGACCAGCCGTTCGATGTGATCAAGCACGTTGACACCCCTGTTCCTGTGGACGTCATCAAGCACGTCCCAGTGC
AGAGGGAGGTGATCCAGCATGTCGAGCAGATCGTCCACAAGCCCGTCCCAGGCCCCGAGCAGATCGTGCAGGTACCCAGGCCCTTCCCAGTGCAT
ATGCAGCCCCCACCACAACCACAGCCTCTTCCCCAGCCTCGTCCAATGATGATGCAGACAACAATGCGAACCCTTCCACCGGTCATGCAGACCCA
ACCCATGCAGGCCCCCCCTCCTCCACAACCGATGCCAACGATGATGACGACCAGCTACATGCCCCCCCCCACCTACTCGCAGCTGCCCACGCAGC
CCTCCTACTCGTACATTGCCCCCCCGGCAACGACTGCCAGTGCCCATGCACCCGTTGCCCCATTCAATTTCCCTCCCCGCCCAGCCGCACCCTTC
GGATTCGGACCCCGCGGCTTTGGCTTCCCTCTACCTGCACCCGGCGTCGGATTTCCCCCACCCAGCAGCCACAGACGATGGCGGCGGCCCACGT
CGGCGCCCCCGTCCCAGGCTCTCCGCCCCCTCCTGCCCCCGCCGTCCAACCTcccgggaccgggtctggtggaactggttcaggtagatct

>gblock (pCvGAPDH–mStayGold–CvTubA1)(2097 bp)
ctgcaggctcagaagATGGTGTCCACCGGGGAGGAGCTCTTCACCGGGGTGGTGCCCTTCAAGTTCCAGCTCAAGGGGACCATCAACGGGAAGTC
CTTCACCGTGGAGGGGGAGGGGGAGGGGAACTCCCACGAGGGGTCCCACAAGGGGAAGTACGTGTGCACCTCCGGGAAGCTCCCCATGTCCTGGG
CCGCCCTCGGGACCTCCTTCGGGTACGGGATGAAGTACTACACCAAGTACCCCTCCGGGCTCAAGAACTGGTTCCACGAGGTGATGCCCGAGGGG
TTCACCTACGACAGACACATCCAGTACAAGGGGGACGGGTCCATCCACGCCAAGCACCAGCACTTCATGAAGAACGGGACCTACCACAACATCGT
GGAGTTCACCGGGCAGGACTTCAAGGAGAACTCCCCCGTGCTCACCGGGGACATGGACGTGTCCCTCCCCAACGAGGTGCAGCACATCCCCATCG
ACGACGGGGTGGAGTGCACCGTGACCCTCCAGTACCCCCTCCTCTCCGACGAGTCCAAGTGCGTGGAGGCCTACCAGAACACCATCATCAAGCCC
CTCCACAACCAGCCCGCCCCCGACGTGCCCTTCCACTGGATCAGAAAGCAGTACACCCAGTCCAAGGACGACACCGAGGAGAGAGACCACATCAT
CCAGTCCGAGACCCTCGAGGCCCACCTCcccgggaccgggtctggtggaactggttcaggtagatctCGCGAGTGCATCTCCATCCACGTCGGAC
AGGCCGGTATCCAGGTCGGAAATGCCTGCTGGGAGCTGTTCTGCCTCGAGCACGGCATTCAGCCTGACGGTCAGATGCCCTCTGACAAGACCATT
GGCGGAGGTGATGATTCTTTCAACACCTTCTTCTCTGAGACCGGTGCCGGCAAGCACGTGCCCCGTTGCGTGATGGTGGATCTTGAGCCCACCGT
CGTCGATGAGGTCCGTACCGGTACCTACCGTCAGCTGTTCCACCCCGAGCAGCTCATCTCCGGAAAGGAGGATGCCGCCAATAACTTCGCCCGTG
GCCACTACACCATCGGAAAGGAGATTGTGGATCTCGTCCTTGACCGTATCCGCAAGCTCGCTGACAACTGCACTGGGCTCCAGGGCTTCCTGATG
TTCAACGCTGTGGGAGGTGGCACAGGCTCCGGTCTTGGCTGCCTTCTCCTCGAGCGTCTGTCTGTCGACTACGGCAAGAAGTCCAAGCTCAACTT
CTGCTCCTGGCCTTCTCCCCAGATCGCCACTGCCGTTGTTGAGCCCTACAACTCCGTGCTCTCCACCCACTCCCTCCTCGAGCACACTGATGTGG
CTGTCATGCTTGATAACGAGGCTCTCTACGATATCTGCCGCAGAAACCTCGACATTGAGCGCCCCACCTACACCAACCTGAACCGTCTGATCGGT
CAGATCATCTCCTCCCTCACCGCCTCCCTCCGTTTCGATGGAGCCCTCAACGTCGACATGACTGAGTTCCAGACCAACTTGGTGCCCTACCCCAG
AATCCACTTCATGCTTTCCTCCTACGCTCCCATCATCTCCGCCGAGAAGGCCTACCACGAGCAGCTCTCTGTCGCAGAGATCACCAACTCTGCCT
TTGAGCCCGCCAACATGATGGCCAAGTGCGACCCCCGCCACGGAAAGTACATGGCCTGCTGCCTCATGTACAGAGGTGACGTCGTCCCCAAGGAC
GTCAACGCAGCCATTGCCACCATCAAGACCAAGCGTACCATCCAGTTCGTGGACTGGTGCCCCACCGGCTTCAAGTGCGGAATCAACTACCAGCC
CCCCACCGTCGTCCCCGGGGGCGATCTTGCTAAGGTGATGCGCGCCTGCTGCATGATCTCCAACTCTACCGCCATCGCTGAGGTCTTCTCCCGCA
TGGACCACAAGTTCGATCTCATGTACGCCAAGCGTGCCTTCGTCCACTGGTACGTCGGTGAGGGTATGGAGGAGGGTGAGTTCTCCGAGGCCCGT
GAGGATCTTGCCGCCCTCGAGAAGGACTACGAGGAGGTCGGCATTGAGACCGCTGAGGCCGAGGGCGAGGAGGAGGGATACGATGAGAACGTCTA
ACTTAAG
