## Supplemental Table 1 for "Transfection of the free-living alga *Chromera velia* enables direct comparisons with its parasitic apicomplexan relative, *Toxoplasma gondii*"

| Name | Sequence |
|---|---|
| P1 | CAAACAAGCAAGCTGCCTTAAGTTACGCCAGAATGCGTTCGCACAG |
| P2 | GTCTCGCTCAGAAGATGAAGAGATCTGTCTTCACACTCGAAGATTTCGTTG |
| P3 | AGAGAGTTTGAGGAGAAGACTGACTCTCGGTCCCTCTTTGATCCG |
| P4 | TCAGTATTTGCGTAAGAGTGCTGCAGCCTCTCACACAAAG |
| P5 | GCCCAGCCAGACGCCTTAAGTTAGGCGAGGATTCTCTCG |
| P6 | CGGATCAAAGAGGGACCGAGAGTCAGTCTTCTCCTCAAACTCTCT |
| Linker | CCCGGGACCGGGTCTGGTGGAACTGGTTCAGGTAGATCT |
